# Structural and functional analysis of a potent sarbecovirus neutralizing antibody

**DOI:** 10.1101/2020.04.07.023903

**Authors:** Dora Pinto, Young-Jun Park, Martina Beltramello, Alexandra C. Walls, M. Alejandra Tortorici, Siro Bianchi, Stefano Jaconi, Katja Culap, Fabrizia Zatta, Anna De Marco, Alessia Peter, Barbara Guarino, Roberto Spreafico, Elisabetta Cameroni, James Brett Case, Rita E. Chen, Colin Havenar-Daughton, Gyorgy Snell, Amalio Telenti, Herbert W. Virgin, Antonio Lanzavecchia, Michael S. Diamond, Katja Fink, David Veesler, Davide Corti

## Abstract

SARS-CoV-2 is a newly emerged coronavirus responsible for the current COVID-19 pandemic that has resulted in more than one million infections and 73,000 deaths^1,2^. Vaccine and therapeutic discovery efforts are paramount to curb the pandemic spread of this zoonotic virus. The SARS-CoV-2 spike (S) glycoprotein promotes entry into host cells and is the main target of neutralizing antibodies. Here we describe multiple monoclonal antibodies targeting SARS-CoV-2 S identified from memory B cells of a SARS survivor infected in 2003. One antibody, named S309, potently neutralizes SARS-CoV-2 and SARS-CoV pseudoviruses as well as authentic SARS-CoV-2 by engaging the S receptor-binding domain. Using cryo-electron microscopy and binding assays, we show that S309 recognizes a glycan-containing epitope that is conserved within the sarbecovirus subgenus, without competing with receptor attachment. Antibody cocktails including S309 along with other antibodies identified here further enhanced SARS-CoV-2 neutralization and may limit the emergence of neutralization-escape mutants. These results pave the way for using S309 and S309-containing antibody cocktails for prophylaxis in individuals at high risk of exposure or as a post-exposure therapy to limit or treat severe disease.

---

Coronavirus entry into host cells is mediated by the transmembrane spike (S) glycoprotein that forms homotrimers protruding from the viral surface^3^. The S glycoprotein comprises two functional subunits: S_1_ (divided into A, B, C and D domains) that is responsible for binding to host cell receptors and S_2_ that promotes fusion of the viral and cellular membranes^4,5^. Both SARS-CoV-2 and SARS-CoV belong to the sarbecovirus subgenus and their S glycoproteins share 80% amino acid sequence identity^6^. SARS-CoV-2 S is closely related to the bat SARS-related CoV (SARSr-CoV) RaTG13 with which it shares 97.2% amino acid sequence identity^1^. We and others recently demonstrated that human angiotensin converting enzyme 2 (hACE2) is a functional receptor for SARS-CoV-2, as is the case for SARS-CoV^1,6-8^. The S domain B (S^B^) is the receptor binding domain (RBD) and binds to hACE2 with high-affinity, possibly contributing to the current rapid SARS-CoV-2 transmission in humans^6,9^, as previously proposed for SARS-CoV^10^.

As the coronavirus S glycoprotein mediates entry into host cells, it is the main target of neutralizing antibodies and the focus of therapeutic and vaccine design efforts^3^. The S trimers are extensively decorated with N-linked glycans that are important for protein folding^11^ and modulate accessibility to host proteases and neutralizing antibodies^12-15^. Cryo-electron microscopy (cryoEM) structures of SARS-CoV-2 S in two distinct functional states^6,9^ along with cryoEM and crystal structures of SARS-CoV-2 S^B^ in complex with hACE2^16-18^ revealed dynamic states of S^B^ domains, providing a blueprint for the design of vaccines and inhibitors of viral entry.

Passive administration of monoclonal antibodies (mAbs) could have a major impact on controlling the SARS-CoV-2 pandemic by providing immediate protection, complementing the development of prophylactic vaccines. Accelerated development of mAbs in a pandemic setting could be reduced to 5-6 months compared to the traditional timeline of 10-12 months (Kelley B., Developing monoclonal antibodies at pandemic speed, Nat Biotechnol, in press). The recent finding that ansuvimab (mAb114) is a safe and effective treatment for symptomatic Ebola virus infection is a striking example of the successful use of mAb therapy during an infectious disease outbreak^19,20^. We previously isolated potently neutralizing human mAbs from memory B cells of individuals infected with SARS-CoV^21^ or MERS-CoV^22^. Passive transfer of these mAbs protected animals challenged with various SARS-CoV isolates and SARS-related CoV (SARSr-CoV)^21,23,24^, as well as with MERS-CoV^22^. Structural characterization of two of these mAbs in complex with SARS-CoV S and MERS-CoV S provided molecular-level information on the mechanisms of viral neutralization^14^. In particular, while both mAbs blocked S^B^ attachment to the host receptor, the SARS-CoV-neutralizing S230 mAb acted by functionally mimicking receptor-attachment and promoting S fusogenic conformational rearrangements^14^. Another mechanism of SARS-CoV neutralization was recently described for mAb CR3022, which bound a cryptic epitope only accessible when at least two out of the three S^B^ domains of a S trimer were in the open conformation^25,26^. However, none of these mAbs neutralize SARS-CoV-2.

## Identification of a potent SARS-CoV-2 neutralizing mAb from a SARS survivor

We previously identified a set of human neutralizing mAbs from an individual infected with SARS-CoV in 2003 that potently inhibited both human and zoonotic SARS-CoV isolates^21,23,27^. To characterize the potential cross-reactivity of these antibodies with SARS-CoV-2, we performed a new memory B cell screening using peripheral blood mononuclear cells collected in 2013 from the same patient. We describe here nineteen mAbs from the initial screen (2004 blood draw)^21,23^ and six mAbs from the new screen (2013 blood draw). The identified mAbs had a broad V gene usage and were not clonally related (**Table 1)**. Eight out of the twenty five mAbs bound to SARS-CoV-2 S and SARS-CoV S transfected CHO cells with EC_50_ values ranging between 1.4 and 6,100 ng/ml, and 0.8 and 254 ng/ml, respectively **(Fig. 1a-b)**. MAbs were further evaluated for binding to the SARS-CoV-2 and SARS-CoV S^B^ domains as well as to the prefusion-stabilized OC43 S^28^, MERS-CoV S^29,30^, SARS-CoV S^30^ and SARS-CoV-2 S^6^ ectodomain trimers. None of the mAbs studied bound to prefusion OC43 S or MERS-CoV S ectodomain trimers, indicating a lack of cross-reactivity outside the sarbecovirus subgenus **(Extended Data Fig.1)**. MAbs S303, S304, S309 and S315 recognized the SARS-CoV-2 and SARS-CoV RBDs. In particular, S309 bound with nanomolar affinity to both S^B^ domains, as determined by biolayer interferometry **(Fig. 1c-d, Extended Data Fig. 2)**. Unexpectedly, S306 and S310 stained cells expressing SARS-CoV-2 S at higher levels than those expressing SARS-CoV S, yet it did not interact with SARS-CoV-2 or SARS-CoV S ectodomain trimers and RBD constructs by ELISA. These results suggest that they may recognize post-fusion SARS-CoV-2 S, which was recently proposed to be abundant on the surface of authentic SARS-CoV-2 viruses^31^ **(Fig. 1a-b and Extended Data Fig.3)**.

**Table 1:**
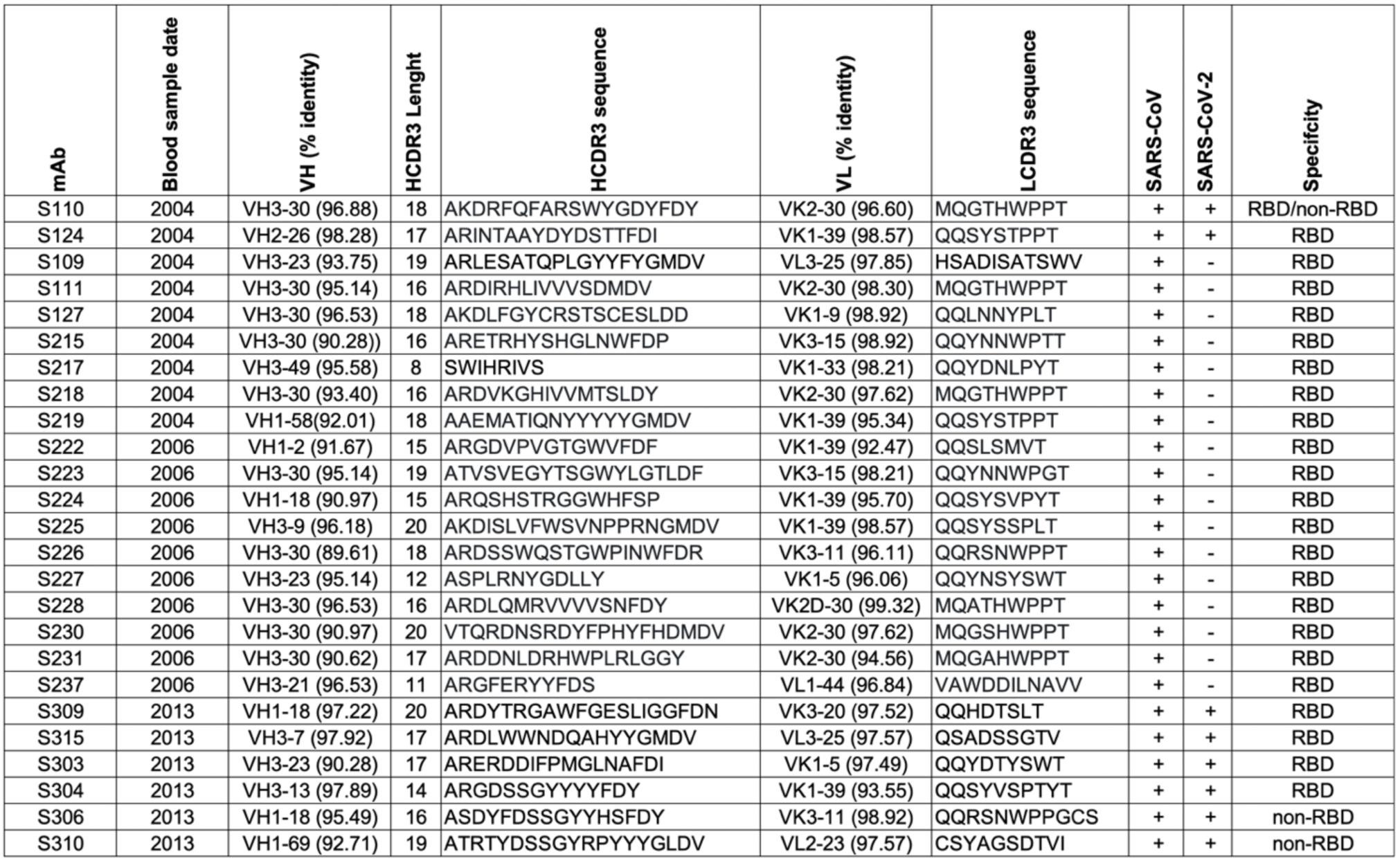
Characteristics of the antibodies described in this study. VH and VL % identity refers to V gene identity compared to germline (IMGT).

**Figure 1:**
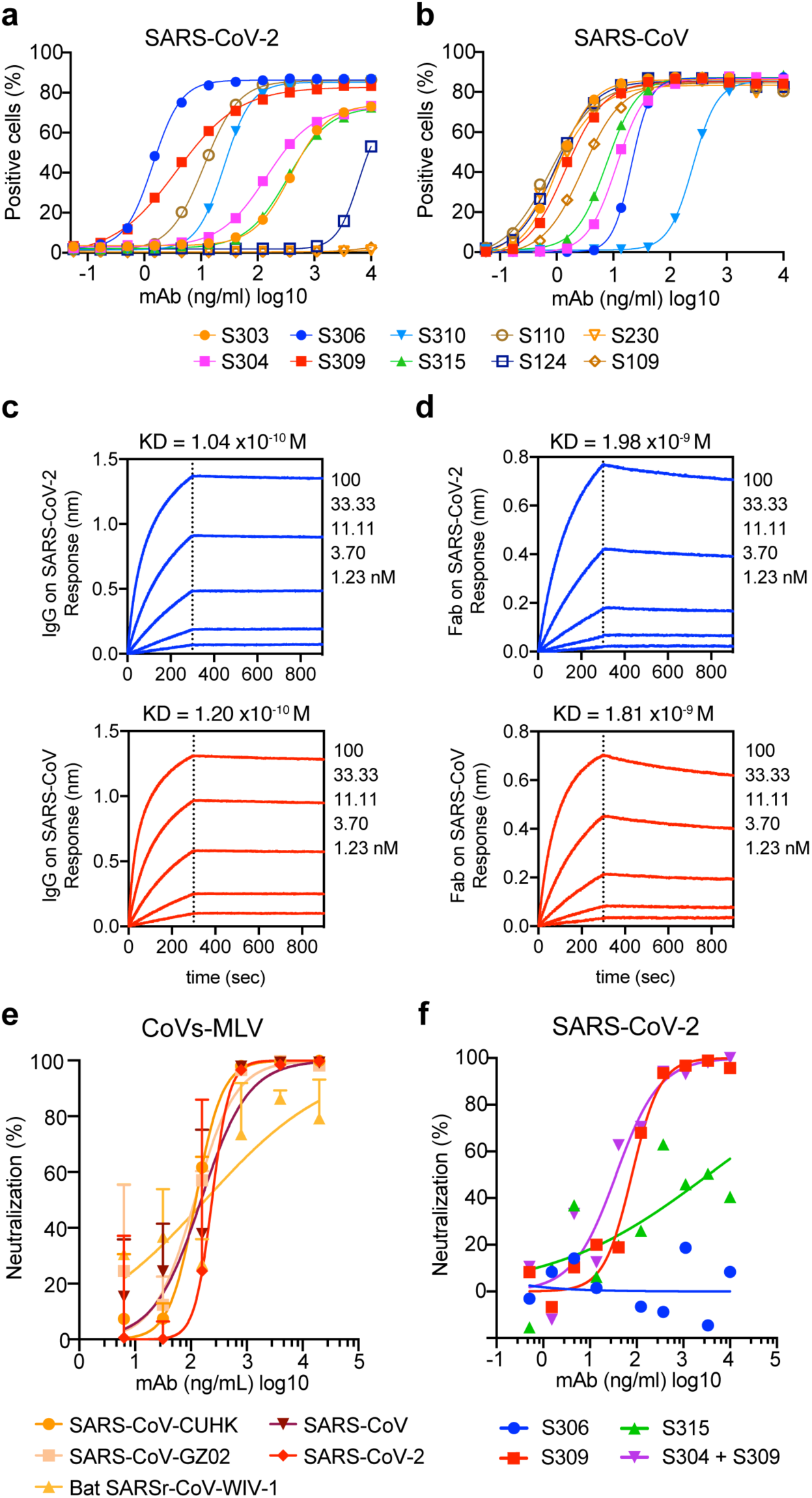
Identification of a potent SARS-CoV-2 neutralizing mAb from a SARS survivor. **a-b**, Binding of a panel of mAbs isolated from a SARS-immune patient to the SARS-CoV-2 (**a**) or SARS-CoV (**b**) S glycoproteins expressed at the surface of ExpiCHO cells (symbols are means of duplicates from one experiment). **c-d**, Affinity measurement of S309 full-length IgG1 and Fab for SARS-CoV-2 and SARS-CoV S^B^ domains measured using biolayer interferometry. **e**, Neutralization of SARS-CoV-2-MLV, SARS-CoV-MLV (bearing S from various isolates) and other sarbecovirus isolates by mAb S309. **f**, Neutralization of authentic SARS-CoV-2 (strain n-CoV/USA_WA1/2020) by mAbs as measured by a focus-forming assay on Vero E6 cells. (**e-f**) mean±SD (**e**) or means (**f)** of duplicates are shown. One representative out of two experiments is shown.

To evaluate the neutralization potency of the SARS-CoV-2 cross-reactive mAbs, we carried out pseudovirus neutralization assays using a murine leukemia virus (MLV) pseudotyping system^32^. S309 showed comparable neutralization potencies against both SARS-CoV and SARS-CoV-2 pseudoviruses, whereas S303 neutralized SARS-CoV-MLV but not SARS-CoV-2-MLV. S304 and S315 weakly neutralized SARS-CoV-MLV and SARS-CoV-2-MLV (**Extended Data Fig.4**). In addition, S309 neutralized SARS-CoV-MLVs from isolates of the 3 phases of the 2002-2003 epidemic with IC_50_ values comprised between 120 and 180 ng/ml and partially neutralized the SARSr-CoV^33^ WIV-1 (**Fig. 1e)**. Finally, mAb S309 potently neutralized authentic SARS-CoV-2 (2019n-CoV/USA_WA1/2020) with an IC_50_ of 69 ng/ml (**Fig. 1f**).

## Structural basis of S309 cross-neutralization of SARS-CoV-2 and SARS-CoV

To study the mechanisms of S309-mediated neutralization, we characterized the complex between the S309 Fab fragment and a prefusion stabilized SARS-CoV-2 S ectodomain trimer^6^ using single-particle cryoEM. Similar to our previous study of apo SARS-CoV-2 S^6^, 3D classification of the cryoEM data enabled identification of two structural states: a trimer with one S^B^ domain open and a closed trimer. We determined 3D reconstructions of the SARS-CoV-2 S ectodomain trimer with a single open S^B^ domain and in a closed state (applying 3-fold symmetry), both with three S309 Fabs bound, at 3.7 Å and 3.3 Å resolution, respectively **(Fig. 2a-c, Extended Data Fig. 5 and Table 2)**. In parallel, we also determined a crystal structure of the S309 Fab at 3.3 Å resolution to assist model building (**Table 3**). The S309 Fab bound to the open S^B^ domain is weakly resolved in the cryoEM map, due to marked conformational variability of the upward pointing S^B^ domain, and was not modeled in density. The analysis below is based on the closed state structure.

**Table 2.**
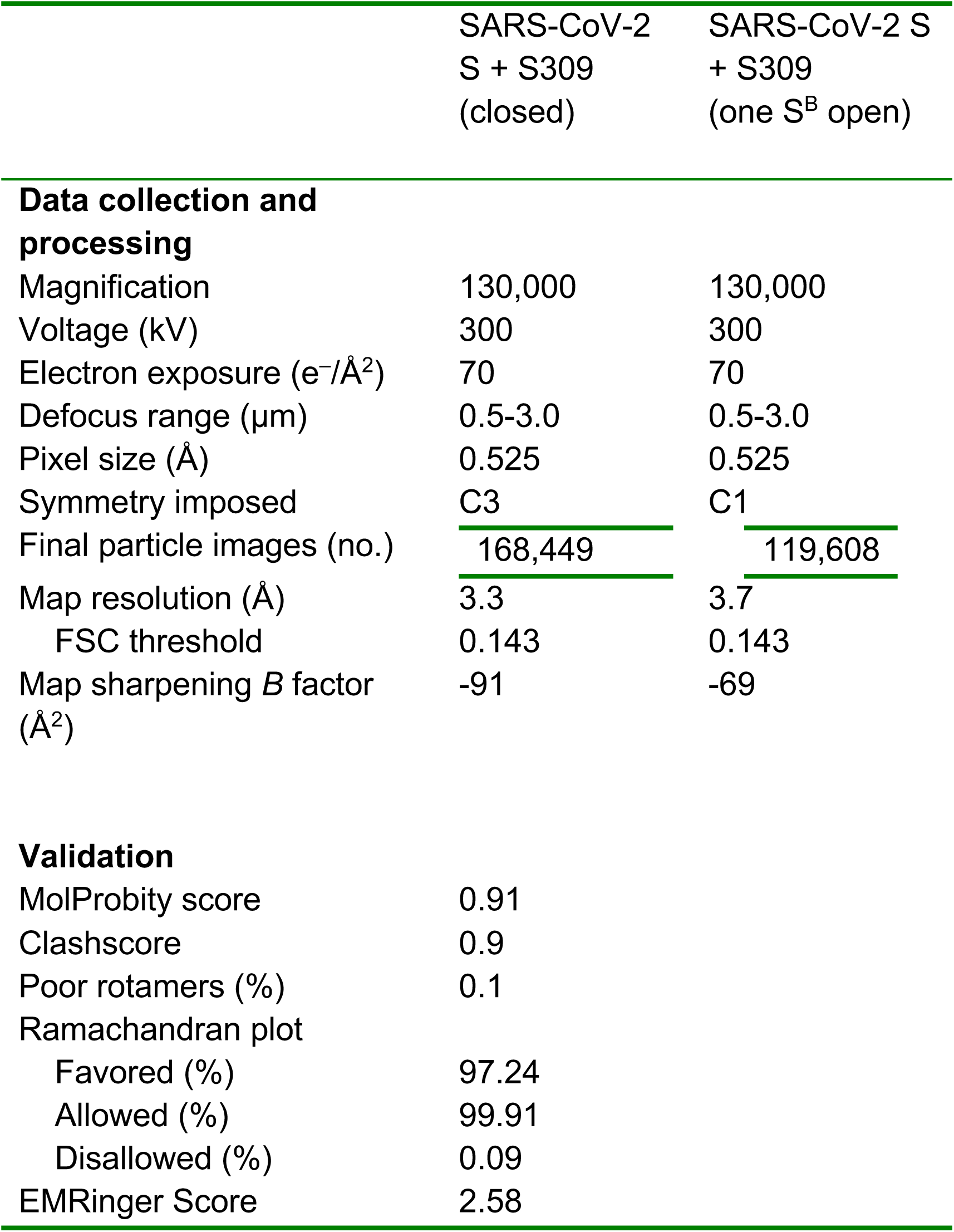
CryoEM data collection and refinement statistics.

**Table 3.** X-ray data collection and refinement statistics.

|  |  |
| --- | --- |
|  | Fab S309 |
| **Data collection** |  |
| Space group | P4<sub>1</sub>2<sub>1</sub>2 |
| Cell constants |  |
| a,b,c (Å) | 132.6, 132.6, 301.2 |
| $\alpha,\beta,\gamma$ (°) | 90, 90, 90 |
| Wavelength (Å) | 0.9812 |
| Resolution (Å) | 68.6 - 3.3 (3.48 - 3.30) |
| Rmerge | 18 (75) |
| I/$\sigma$(I) | 13.2 (2) |
| CC(1/2) | 99.0 (33) |
| Completeness (%) | 99.4 (99.0) |
| Redundancy | 12 |
| **Refinement** |  |
| Resolution (Å) | 68.6 - 3.3 |
| Unique reflections | 41,395 |
| Rwork/Rfree (%) | 20.5 / 23.8 |
| Number of protein atoms | 8,347 |
| Number of water atoms | 0 |
| R.m.s.d. bond lengths (Å) | 0.06 |
| R.m.s.d. bond angles (°) | 1.45 |
| Favored Ramachandran residues (%) | 96 |
| Allowed Ramachandran residues (%) | 3.54 |
| Disallowed Ramachandran residues (%) | 0.46 |

<sup>1</sup>Numbers in parentheses refer to outer resolution shell

**Figure 2:**
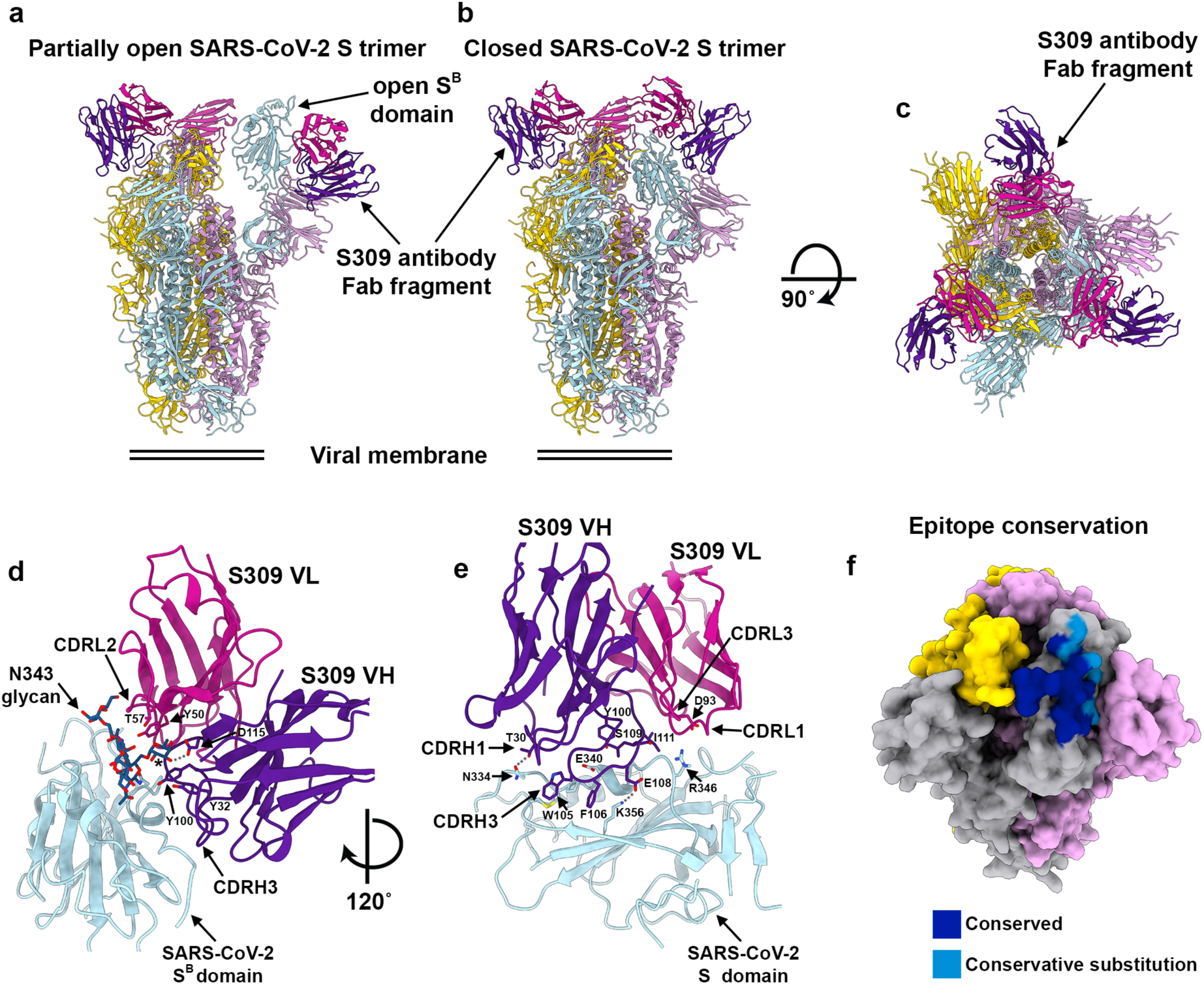
CryoEM structures of the SARS-CoV-2 S glycoprotein in complex with the S309 neutralizing antibody Fab fragment. **a**, Ribbon diagram of the partially open SARS-CoV-2 S trimer (one S^B^ domain is open) bound to three S309 Fabs. **b-c**, Ribbon diagrams of the closed SARS-CoV-2 S trimer bound to three S309 Fabs shown in two orthogonal orientations. **d**, Close-up view of the S309 epitope showing the contacts formed with the core fucose (labeled with a star) and the core N-acetyl-glucosamine of the oligosaccharide at position N343. **e**, Close-up view of the S309 epitope showing the 20-residue long CDRH3 siting atop the S^B^ helix comprising residues 337-344. The oligosaccharide at position N343 is omitted for clarity. In panels (**c-d**), selected residues involved in interactions between S309 and SARS-CoV-2 S are shown. **F**, Molecular surface representation of the SARS-CoV-2 S trimer showing the S309 footprint colored by residue conservation on one protomer among SARS-CoV-2 and SARS-CoV S glycoproteins. The other two protomers are colored pink and gold.

S309 recognizes a protein/glycan epitope on the SARS-CoV-2 S^B^, distinct from the receptor-binding motif. The epitope is accessible in both the open and closed S states, explaining the stoichiometric binding of Fab to the S trimer **(Fig. 2a-c)**. The S309 paratope is composed of all six CDR loops that burie a surface area of ∼1,050Å^2^ at the interface with S^B^ through electrostatic interactions and hydrophobic contacts. The 20-residue long CDRH3 sits atop the S^B^ helix comprising residues 337-344 and also contacts the edge of the S^B^ five-stranded β-sheet (residues 356-361), overall accounting for ∼50% of the buried surface area **(Fig. 2d-e)**. CDRL1 and CDRL2 extend the epitope by interacting with the helix spanning residues 440-444 that is located near the S 3-fold molecular axis. CDRH3 and CDRL2 sandwich the SARS-CoV-2 S glycan at position N343 through contacts with the core fucose moiety (in agreement with the detection of SARS-CoV-2 N343 core-fucosylated peptides by mass-spectrometry^34^) and to a lesser extent with the core N-acetyl-glucosamine (Fig. 2d). These latter interactions bury an average surface of ∼170 Å^2^ and stabilize the N343 oligosaccharide which is resolved to a much larger extent than in the apo SARS-CoV-2 S structures^6,9^.

The structural data explain the S309 cross-reactivity between SARS-CoV-2 and SARS-CoV as 19 out of 24 residues of the epitope are strictly conserved (**Fig. 2f** and **Extended Data Fig. 6a-b)**. R346_SARS-CoV-2_, R357_SARS-CoV-2_, N354_SARS-CoV-2_ and L441_SARS-CoV-2_ are conservatively substituted to K333_SARS-CoV_, K344_SARS-CoV_ (except for SARS-CoV isolate GZ02 where it is R444_SARS-CoV_), E341_SARS-CoV_ and I428_SARS-CoV_ whereas K444_SARS-CoV-2_ is semi-conservatively substituted to T431_SARS-CoV_, in agreement with the comparable binding affinities to SARS-CoV and SARS-CoV-2 S (**Fig. 1c)**. The oligosaccharide at position N343 is also conserved in both viruses and corresponds to SARS-CoV N330, for which we previously detected core-fucosylated glycopeptides by mass spectrometry^14^ which would allow for similar interactions with the S309 Fab. Analysis of the S glycoprotein sequences of the 2,229 SARS-CoV-2 isolates reported to date indicates that several mutations have occurred with variable frequency on the SARS-CoV-2 S ectodomain (**Extended Data Fig. 7a-b**) but no mutations arose within the epitope recognized by S309 mAb. Finally, S309 contact residues showed a high degree of conservation across clade 1, 2 and 3 sarbecovirus human and animal isolates^35^ (**Extended Data Fig. 7c)**. Collectively, the structural data indicate that S309 could neutralize all SARS-CoV-2 isolates circulating to date and possibly most other zoonotic sarbecoviruses.

## Mechanism of S309-mediated neutralization of SARS-CoV-2 and SARS-CoV

The cryoEM structure of S309 bound to SARS-CoV-2 S presented here combined with the structures of SARS-CoV-2 S^B^ and SARS-CoV S^B^ in complex with ACE2^16-18,36^ indicate that the Fab engages an epitope distinct from the receptor-binding motif and would not clash with ACE2 upon binding to S **(Figure 3a-b)**. Biolayer interferometry analysis of S309 Fab or IgG binding to the SARS-CoV-2 S^B^ domain or the S ectodomain trimer confirmed the absence of competition between the mAb and ACE2 for binding to SARS-CoV-2 S **(Figure 3c and Extended Data Fig. 8)**.

**Figure 3:**
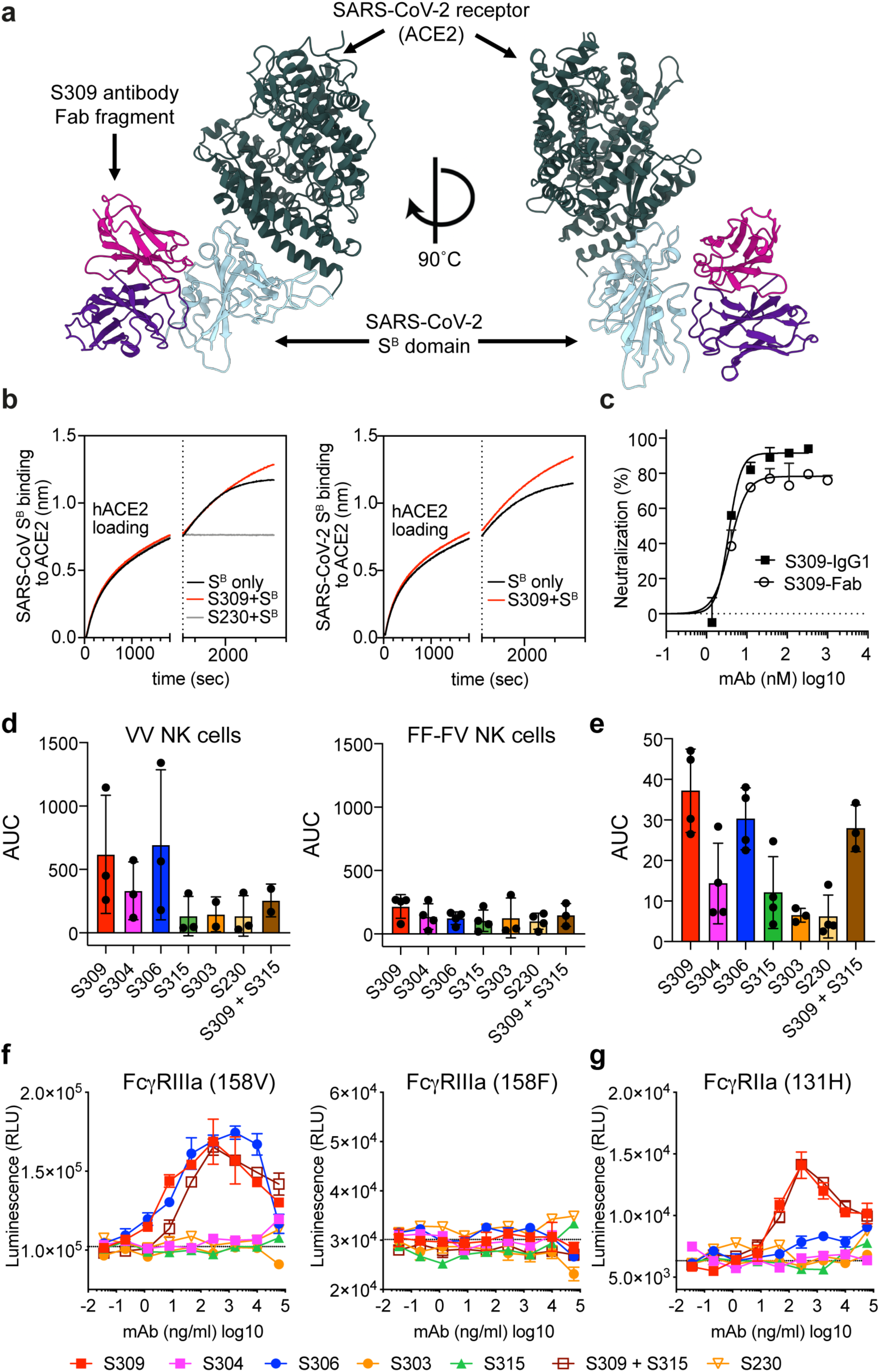
Mechanism of neutralization of S309 mAb. **a-b**, Ribbon diagrams of S309 and ACE2 bound to SARS-CoV-2 S^B^. This composite model was generated using the SARS-CoV-2 S/S309 cryoEM structure reported here and a crystal structure of SARS-CoV-2 S bound to ACE2^16^. **c**, Competition of S309 or S230 mAbs with ACE2 to bind to SARS-CoV S^B^ (left panel) and SARS-CoV-2 S^B^ (right panel). ACE2 was immobilized at the surface of biosensors before incubation with S^B^ domain alone or S^B^ precomplexed with mAbs. The vertical dashed line indicates the start of the association of mAb-complexed or free S^B^ to solid-phase ACE2. **d**, Neutralization of SARS-CoV-MLV by S309 IgG1 or S309 Fab, plotted in nM (means <u>±</u>SD is shown, one out of two experiments is shown). **e**, mAb-mediated ADCC using primary NK effector cells and SARS-CoV-2 S-expressing ExpiCHO as target cells. Bar graph shows the average area under the curve (AUC) for the responses of 3-4 donors genotyped for their FcγRIIIa (mean±SD, from two independent experiments). **f**, Activation of high affinity (V158) or low affinity (F158) FcγRIIIa was measured using Jurkat reporter cells and SARS-CoV-2 S-expressing ExpiCHO as target cells (one experiment, one or two measurements per mAb). **g**, mAb-mediated ADCP using Cell Trace Violet-labelled PBMCs as phagocytic cells and PKF67-labelled SARS-CoV-2 S-expressing ExpiCHO as target cells. Bar graph shows the average area under the curve (AUC) for the responses of four donor (mean±SD, from two independent experiments). **h**, Activation of FcγRIIa measured using Jurkat reporter cells and SARS-CoV-2 S-expressing ExpiCHO as target cells (one experiment, one or two measurements per mAb).

To further investigate the mechanism of S309-mediated neutralization, we compared side-by-side transduction of SARS-CoV-2-MLV in the presence of either S309 Fab or S309 IgG. Both experiments yielded comparable IC_50_ values (3.8 and 3.5 nM, respectively), indicating similar potencies for IgG and Fab **(Fig. 3d)**. However, The S309 IgG reached 100% neutralization, whereas the S309 Fab plateaued at ∼80% neutralization **(Fig. 3d)**. This result indicates that one or more IgG-specific bivalent mechanisms, such as S trimer cross-linking, steric hindrance or aggregation of virions^37^, may contribute to the ability to fully neutralize pseudovirions.

Fc-dependent effector mechanisms, such as NK-mediated antibody-dependent cell cytotoxicity (ADCC) and antibody-dependent cellular phagocytosis (ADCP) can contribute to viral control in infected individuals. We observed efficient S309- and S306-mediated ADCC of SARS-CoV-2 S-transfected cells, whereas the other mAbs tested showed limited or no activity (**Fig. 3e and Extended Data Fig. 9a**). These findings might be related to distinct binding orientations and/or positioning of the mAb Fc fragment relative to the FcγRIIIa receptors. ADCC was observed only using NK (effector) cells expressing the high-affinity FcγRIIIa variant (V158) but not the low-affinity variant (F158) (**Fig. 3e**). These results, which we confirmed using a FcγRIIIa cell reporter assay (**Fig. 3f**), suggest that S309 Fc engineering could potentially enhance activation of NK cells with the low-affinity FcγRIIIa variant (F158)^38^. Macrophage or dendritic cell-mediated ADCP can contribute to viral control by clearing virus and infected cells and by stimulating T cell response via presentation of viral antigens^39,40^. Similar to the ADCC results, mAbs S309 and S306 showed the strongest ADCP response **(Fig. 3g** and **Extended Data Fig. 8b)**. FcγRIIa signaling, however, was only observed for S309 (**Fig. 3h**). These findings suggest that ADCP by monocytes was dependent on both FcγRIIIa and FcγRIIa engagement. Collectively, these results demonstrate that in addition to potent *in vitro* neutralization, S309 may leverage additional protective mechanisms *in vivo*, as previously shown for other antiviral antibodies^41,42^.

## MAb cocktails enhance SARS-CoV-2 neutralization

To gain more insight into the epitopes recognized by our panel of mAbs, we used structural information, escape mutants analysis ^23,27,30^, and biolayer inteferometry-based epitope binning to map the antigenic sites present on the SARS-CoV and SARS-CoV-2 S^B^ domains (**Fig. 4a and Extended Data Fig.10)**. This analysis identified at least four antigenic sites within the S^B^ domain of SARS-CoV targeted by our panel of mAbs. The receptor-binding motif, which is targeted by S230, S227 and S110, is termed site I. Sites II and III are defined by S315 and S124, respectively, and the two sites were bridged by mAb S304. Site IV is defined by S309, S109, and S303 mAbs. Given the lower number of mAbs cross-reacting with SARS-CoV-2, we were able to identify sites IV targeted by S309 and S303, and site II-III targeted by S304 and S315 (**Fig. 4b**).

**Figure 4:**
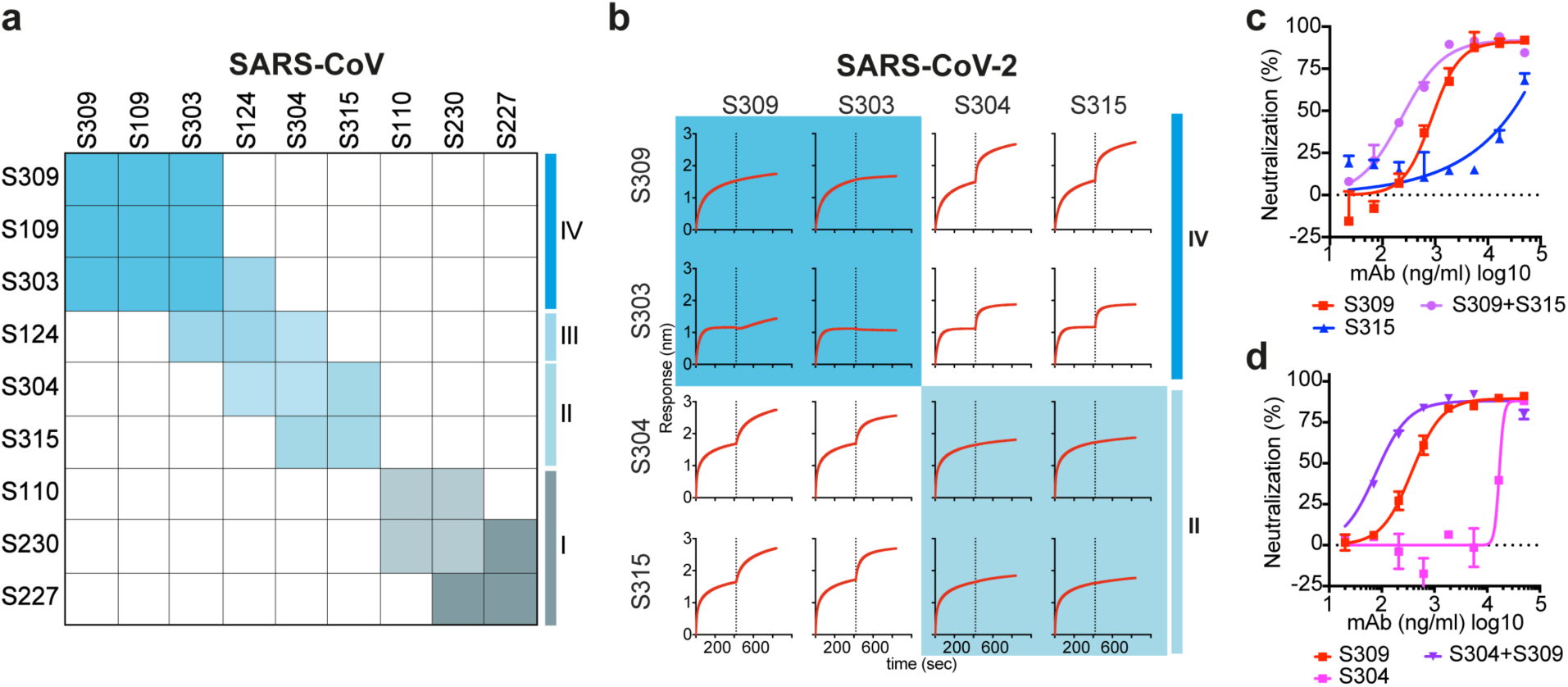
MAb cocktails enhance SARS-CoV-2 neutralization. a, Heat map showing the competition of mAb pairs for binding to the SARS-CoV S^B^ domain as measured by biolayer interferometry (as shown in Extended Data Fig. 9). **b**, Competition of mAb pairs for binding to the SARS-CoV-2 S^B^ domain. **c-d**, Neutralization of SARS-CoV-2-MLV by S309 combined with an equimolar amount of S304 or S315 mAbs. For mAb cocktails the concentration on the x axis is that of the individual mAbs.

Based on the above findings, we evaluated the neutralization potency of the site IV S309 mAb in combination with either the site II S315 mAb or site II-III S304 mAb. Although S304 and S315 alone were weakly neutralizing, the combination of either of these mAbs with S309 resulted in an enhanced neutralization potency, compared to single mAbs, against both SARS-CoV-2-MLV and authentic SARS-CoV-2 **(Fig. 4c-d** and **Fig. 1e)**. A synergistic effect between two non-competing anti-RBD mAbs has been already reported for SARS-CoV^43^ and our data extend this observation to SARS-CoV-2, providing a proof-of-concept for the use of mAbs combinations to prevent or control SARS-CoV-2.

In summary, our study identifies S309 as a human mAb with broad neutralizing activity against multiple sarbecoviruses, including SARS-CoV-2, via recognition of a highly conserved epitope in the S^B^ domain comprising the N343-glycan (N330 in SARS-CoV S). Furthermore, S309 can recruit effector mechanisms and synergizes with weakly neutralizing mAbs, which may mitigate the risk of viral escape. Our data indicate the potential to discover potently neutralizing pan-sarbecovirus mAbs, highlight antigenic sites to include in vaccine design, and pave the way to support preparedness for future sarbecovirus outbreaks. As S309 bears the promise to be an effective countermeasure to curtail the COVID-19 pandemic caused by SARS-CoV-2, Fc variants of S309 with increased half-life and effector functions have entered an accelerated development path towards clinical trials.

## ACKNOWLEDGEMENTS

This study was supported by the National Institute of General Medical Sciences (R01GM120553, D.V.), the National Institute of Allergy and Infectious Diseases (HHSN272201700059C to DV and 75N93019C00062 to MSD)), a Pew Biomedical Scholars Award (D.V.), an Investigators in the Pathogenesis of Infectious Disease Award from the Burroughs Wellcome Fund (D.V.), the University of Washington Arnold and Mabel Beckman cryoEM center, the Pasteur Institute (M.A.T.) and the beamline 5.0.1 at the Advanced Light Source at Lawrence Berkley National Laboratory. We gratefully acknowledge the authors, originating and submitting laboratories of the sequences from GISAID’s EpiFlu™ Database upon which this research is based.

## AUTHOR CONTRIBUTIONS

A.C.W., K.F., M.S.D., D.V. and D.C. designed the experiments. A.C.W., M.A.T., S.J., E.C. expressed and purified the proteins. K.C., F.Z., S.J., E.C. sequenced and expressed antibodies. D.P., M.B., A.C.W. and S.B. performed binding assays. D.P., M.B., A.C.W., A.P., A.D.M. carried out pseudovirus neutralization assays. J.B.C., R.E.C. performed neutralization assays with authentic SARS-CoV-2. B.G. performed effector function assays. Y.J.P. prepared samples for cryoEM and collected the data. Y.J.P. and D.V. processed the data, built and refined the atomic models. A.C.W. crystallized the S309 Fab. Y.J.P. collected and processed the X-ray diffraction data and built and refined the atomic model. R.S., A.T. and G.S. performed bioinformatic and conservation analysis. A.L. provided key reagents. A.C.W., K.F., C.H.D., H.W.V., A.L., D.V., D.C. analyzed the data and prepared the manuscript with input from all authors.

## MATERIALS AND METHODS

### Ethics statement

Donors provided written informed consent for the use of blood and blood components (such as sera), following approval by the Canton Ticino Ethics Committee, Switzerland.

### Antibody discovery and expression

Monoclonal antibodies were isolated from EBV-immortalized memory B cells. Recombinant antibodies were expressed in ExpiCHO cells transiently co-transfected with plasmids expressing the heavy and light chain, as previously described^44^. Abs S303, S304, S306, S309, S310 and S315 were expressed as rIgG-LS antibodies. The LS mutation confers a longer half-life in vivo^45^. Antibodies S110 and S124 tested in Fig. 1 and Extended Data Fig. 1 were purified mAbs produced from immortalized B cells.

### Transient expression of recombinant SARS-CoV-2 protein and flow cytometry

The full-length S gene of SARS-CoV-2 strain (SARS-CoV-2-S) isolate BetaCoV/Wuhan-Hu-1/2019 (accession number MN908947) was codon optimized for human cell expression and cloned into the phCMV1 expression vector (Genlantis). Expi-CHO cells were transiently transfected with phCMV1-SARS-CoV-2-S, SARS-spike_pcDNA.3 (strain SARS) or empty phCMV1 (Mock) using Expifectamine CHO Enhancer. Two days after transfection, cells were collected for immunostaining with mAbs. An Alexa647-labelled secondary antibody anti-human IgG Fc was used for detection. Binding of mAbs to transfected cells was analyzed by flow-cytometry using a ZE5 Cell Analyzer (Biorard) and FlowJo software (TreeStar). Positive binding was defined by differential staining of CoV-S-transfectants versus mock-transfectants.

### Affinity determination and competition experiments using Octet (BLI, biolayer interferometry)

KD determination of full-length antibodies: Protein A biosensors (Pall ForteBio) were used to immobilize recombinant antibodies at 2.7 μg/ml for 1min, after a hydration step for 10 min with Kinetics Buffer (KB; 0.01% endotoxin-free BSA, 0.002% Tween-20, 0.005% NaN3 in PBS). Association curves were recorded for 5 minutes by incubating the mAb-coated sensors with different concentration of SARS-CoV RBD (Sino Biological) or SARS-CoV-2 RBD (produced in house; residues 331-550 of spike protein from BetaCoV/Wuhan-Hu-1/2019, accession number MN908947). The highest RBD concentration was 10 μg/ml, then serially diluted 1:2.5. Dissociation was recorded for 9 minutes by moving the sensors to wells containing KB. KD values were calculated using a global fit model (Octet). Octet Red96 (ForteBio) equipment was used.

KD determination of full-length antibodies compared to Fab: His-tagged RBD of SARS-CoV or SARS-CoV-2 were loaded at 3 µg/ml in KB for 15 minutes onto anti-HIS (HIS2) biosensors (Molecular Devices, ForteBio). Association of mAb and Fab was performed in KB at 15 μg/ml and 5 μg/ml respectively for 5 minutes. Dissociation in KB was measured for 10 minutes.

MAbs competition experiments: His-tagged RBD of SARS-CoV or SARS-CoV-2 was loaded for 5 minutes at 3 μg/ml in KB onto anti-Penta-HIS (HIS1K) biosensors (Molecular Devices, ForteBio). Association of mAbs was performed in KB at 15 μg/ml.

ACE2 competition experiments: ACE2-His (Bio-Techne AG) was loaded for 30 minutes at 5 µg/ml in KB onto anti-HIS (HIS2) biosensors (Molecular Devices-ForteBio).

SARS-CoV RBD-rabbitFc or SARS-CoV-2 RBD-mouseFc (Sino Biological Europe GmbH) at 1 µg/ml was associated for 15 minutes, after a preincubation with or without Ab (30 µg/ml, 30 minutes). Dissociation was monitored for 5 minutes.

### ELISA

The following proteins were coated on 96 well ELISA plates at the following concentrations: SARS-CoV RBD (Sino Biological, 40150-V08B1) at 1 µg/ml, SARS-CoV-2 RBD (produced in house) at 10 µg/ml, ectodomains (stabilized prefusion trimer) of SARS-CoV, SARS-CoV-2, OC43 and MERS, all at 1ug/ml. After blocking with 1% BSA in PBS, antibodies es were added to the plates in a concentration range between 5 and 0.000028 μg/ml and incubated for 1 h at RT. Plates were washed and secondary Ab Goat Anti Human IgG-AP (Southern Biotechnology: 2040-04) was added. Substrate P-NitroPhenyl Phosphate (pNPP) (Sigma-Aldrich 71768) was used for colour development. OD405 was read on an ELx808IU plate reader (Biotek).

### Measurement of Fc-effector functions

ADCC assays were performed using ExpiCHO-S cells transient transfected with SARS-CoV or SARS-CoV-2 S as targets. Target cells were incubated with titrated concentrations of mAbs and after 10 minutes incubated with primary human NK cells as effector cells at an effector:target ratio of 9:1. NK cells were isolated from fresh blood of healthy donors using the MACSxpress NK Isolation Kit (Miltenyi Biotec, Cat. Nr.: 130-098-185). ADCC was measured using LDH release assay (Cytotoxicity Detection Kit (LDH) (Roche; Cat. Nr.: 11644793001) after 4 hours of incubation at 37°C.

ADCP assays were performed using ExpiCHO-S target cells transiently transfected with SARS-CoV-2 S and fluorescently labeled with PKH67 Fluorescent Cell Linker Kits (Sigma Aldrich, Cat. Nr.: MINI67) as targets. Target cells were incubated with titrated concentrations of mAbs for 10minutes, followed by incubation with human PBMCs isolated from healthy donors that were fluorescently labeled with Cell Trace Violet (Invitrogen, Cat. Nr.: C34557) at an effector:target ratio of 20:1. After an overnight incubation at 37°C, cells were stained with anti-human CD14-APC antibody (BD Pharmingen, Cat. Nr.: 561708, Clone M5E2) to stain monocytes. Antibody-mediated phagocytosis was determined by flow cytometry, gating on CD14^+^ cells that were double positive for cell trace violet and PKH67.

Determination of mAb-dependent activation of human FcγRIIIa or FcγRIIa was performed using ExpiCHO cells transiently transfected with SARS-CoV-2 S (BetaCoV/Wuhan-Hu-1/2019), incubated with titrated concentrations of mAbs for 10 minutes. ExpiCHO cells then were incubated with Jurkat cells expressing FcγRIIIa receptor or FcγRIIa on their surface and stably transfected with NFAT-driven luciferase gene (Promega, Cat. Nr.: G9798 and G7018) at an effector to target ratio of 6:1 for FcγRIIIa and 5:1 for FcγRIIa. Activation of human FcγRs in this bioassay results in the NFAT-mediated expression of the luciferase reporter gene. Luminescence was measured after 21 hours of incubation at 37°C with 5% CO_2,_ using the Bio-Glo-TM Luciferase Assay Reagent according to the manufacturer’s instructions.

### Pseudovirus neutralization assays

Murine leukemia virus (MLV)-based SARS-CoV S-pseudotyped viruses were prepared as previously described^6,32^. HEK293T cells were co-transfected with a SARS-CoV, SARS-CoV-2, CUHK, GZ02, or WiV1 S encoding-plasmid, an MLV Gag-Pol packaging construct and the MLV transfer vector encoding aluciferase reporter using the Lipofectamine 2000 transfection reagent (Life Technologies) according to the manufacturer’s instructions. Cells were incubated for 5 hours at 37°C with 8% CO_2_ with OPTIMEM transfection medium. DMEM containing 10% FBS was added for 72 hours. VeroE6 cells or DBT cells transfected with human ACE2 were cultured in DMEM containing 10% FBS, 1% PenStrep and plated into 96 well plates for 16-24 hours. Concentrated pseudovirus with or without serial dilution of antibodies was incubated for 1 hour and then added to the wells after washing 3X with DMEM. After 2-3 hours DMEM containing 20% FBS and 2% PenStrep was added to the cells for 48 hours. Following 48 hours of infection, One-Glo-EX (Promega) was added to the cells and incubated in the dark for 5-10 minutes prior to reading on a Varioskan LUX plate reader (ThermoFisher). Measurements were done in duplicate and relative luciferase units (RLU) were converted to percent neutralization and plotted with a non-linear regression curve fit in PRISM.

### Live virus neutralization assay

SARS-CoV-2 strain 2019-nCoV/USA_WA1/2020 was obtained from the Centers for Disease Control and Prevention (gift of Natalie Thornburg). Virus was passaged once in Vero CCL81 cells (ATCC) and titrated by focus-forming assay on Vero E6 cells. Serial dilutions of indicated mAbs were incubated with 10^2^ focus forming units (FFU) of SARS-CoV-2 for 1 hour at 37°C. MAb-virus complexes were added to Vero E6 cell monolayers in 96-well plates and incubated at 37°C for 1 hour. Subsequently, cells were overlaid with 1% (w/v) methylcellulose in MEM supplemented with 2% FBS. Plates were harvested 30 hours later by removing overlays and fixed with 4% PFA in PBS for 20 minutes at room temperature. Plates were washed and sequentially incubated with 1 µg/mL of CR3022^46^ anti-S antibody and HRP-conjugated goat anti-human IgG in PBS supplemented with 0.1% saponin and 0.1% BSA. SARS-CoV-2-infected cell foci were visualized using TrueBlue peroxidase substrate (KPL) and quantitated on an ImmunoSpot microanalyzer (Cellular Technologies). Data were processed using Prism software (GraphPad Prism 8.0).

### Recombinant Spike ectodomain production

The SARS-CoV-2 2P S (Genbank: YP_009724390.1) ectodomain was produced in 500mL cultures of HEK293F cells grown in suspension using FreeStyle 293 expression medium (Life technologies) at 37°C in a humidified 8% CO2 incubator rotating at 130 r.p.m, as previously reported^6^. The culture was transfected using 293fectin (ThermoFisher Scientific) with cells grown to a density of 10^6^ cells per mL and cultivated for three days. The supernatant was harvested and cells were resuspended for another three days, yielding two harvests. Clarified supernatants were purified using a 5mL Cobalt affinity column (Takara). Purified protein was filtered or concentrated and flash frozen in a buffer containing 50 mM Tris pH 8.0 and 150 mM NaCl prior to cryoEM analysis. The SARS-CoV S, HCoV-OC43 S and MERS-CoV S constructs were previously described^14,28^ and produced similarly to SARS-CoV-2 2P S.

### CryoEM sample preparation and data collection

3 µL of SARS-CoV-2 S at 1.6 mg/mL was mixed with 0.45 µL of S309 Fab at 7.4 mg/mL for 1 min at room temperature before application onto a freshly glow discharged 1.2/1.3 UltraFoil grid (300 mesh). Plunge freezing used a vitrobot MarkIV (ThermoFisher Scientific) using a blot force of 0 and 6.5 second blot time at 100% humidity and 25°C. Data were acquired using the Leginon software ^47^ to control an FEI Titan Krios transmission electron microscope operated at 300 kV and equipped with a Gatan K2 Summit direct detector and Gatan Quantum GIF energy filter, operated in zero-loss mode with a slit width of 20 eV. Automated data collection was carried out using Leginon at a nominal magnification of 130,000x with a pixel size of 0.525 Å with tilt angles ranging between 20° and 50°, as previously described^48^. The dose rate was adjusted to 8 counts/pixel/s, and each movie was acquired in super-resolution mode fractionated in 50 frames of 200 ms. 3,900 micrographs were collected in a single session with a defocus range comprised between 1.0 and 3.0 μm.

### CryoEM data processing

Movie frame alignment, estimation of the microscope contrast-transfer function parameters, particle picking and extraction were carried out using Warp ^49^. Particle images were extracted with a box size of 800 binned to 400 yielding a pixel size of 1.05 Å. For each data set two rounds of reference-free 2D classification were performed using cryoSPARC ^50^ to select well-defined particle images. Subsequently, two rounds of 3D classification with 50 iterations each (angular sampling 7.5° for 25 iterations and 1.8° with local search for 25 iterations), using our previously reported closed SARS-CoV-2 S structure^6^ as initial model, were carried out using Relion ^51^ without imposing symmetry to separate distinct SARS-CoV-2 S conformations. 3D refinements were carried out using non-uniform refinement along with per-particle defocus refinement in cryoSPARC^50^. Particle images were subjected to Bayesian polishing ^52^ before performing another round of non-uniform refinement in cryoSPARC ^50^ followed by per-particle defocus refinement and again non-uniform refinement. Reported resolutions are based on the gold-standard Fourier shell correlation (FSC) of 0.143 criterion and Fourier shell correlation curves were corrected for the effects of soft masking by high-resolution noise substitution^53^.

### CryoEM model building and analysis

UCSF Chimera ^54^ and Coot were used to fit atomic models (PDB 6VXX and PDB 6VYB) into the cryoEM maps. The Fab was subsequently manually built using Coot^55,56^. N-linked glycans were hand-built into the density where visible and the models were refined and relaxed using Rosetta^57^. Glycan refinement relied on a dedicated Rosetta protocol, which uses physically realistic geometries based on prior knowledge of saccharide chemical properties ^58^, and was aided by using both sharpened and unsharpened maps. Models were analyzed using MolProbity ^59^, EMringer ^60^, Phenix ^61^ and privateer ^62^ to validate the stereochemistry of both the protein and glycan components. Figures were generated using UCSF ChimeraX ^63^.

### Crystallization and X-ray structure determination of Fab S309

Fab S309 crystals were grown in hanging drop set up with a mosquito at 20°C using 150 nL protein solution in Tris HCl pH 8.0, 150 mM NaCl and 150nL mother liquor solution containing 1.1 M Sodium Malonate, 0.1 M HEPES, pH 7.0 and 0.5% (w/v) Jeffamine ED-2001. Crystals were cryo-protected using the mother liquor solution supplemented with 30% glycerol. The dataset was collected at ALS beamline 5.0.2 and processed to 3.3 Å resolution in space group P4_1_2_1_2 using mosflm^64^ and Aimless^65^. The structure of Fab S309 was solved by molecular replacement using Phaser^66^ and homology models as search models. The coordinates were improved and completed using Coot^55^ and refined with REFMAC5^67^. Crystallographic data collection and refinement statistics are shown in Table 3.

### Sequence alignment

SARS-CoV-2 genomics sequences were downloaded from GISAID on March 29^th^ 2020, using the “complete (>29,000 bp)” and “low coverage exclusion” filters. Bat and pangolin sequences were removed to yield human-only sequences. The spike ORF was localized by performing reference protein (YP_009724390.1)-genome alignments with GeneWise2. Incomplete matches and indel-containing ORFs were rescued and included in downstream analysis. Nucleotide sequences were translated *in silico* using seqkit. Sequences with more than 10% undetermined aminoacids (due to N basecalls) were removed. Multiple sequence alignment was performed using MAFFT. Variants were determined by comparison of aligned sequences (n=2,229) to the reference sequence using the R/Bioconductor package Biostrings. A similar strategy was used to extract and translate spike protein sequences from SARS-CoV genomes sourced from ViPR (search criteria: SARS-related coronavirus, full-length genomes, human host, deposited before December 2019 to exclude SARS-CoV-2, n=53). Sourced SARS-CoV genome sequences comprised all the major published strains, such as Urbani, Tor2, TW1, P2, Frankfurt1, among others. Pangolin sequences as shown by Tsan-Yuk Lam et al^68^ were sourced from GISAID. Bat sequences from the three clades of sarbecoviruses as shown by Lu et al^35^ were sourced from Genbank. Civet and racoon dog sequences were similarly sourced from Genbank.

