## Supplementary Figures for "Structural and functional analysis of a potent sarbecovirus neutralizing antibody"

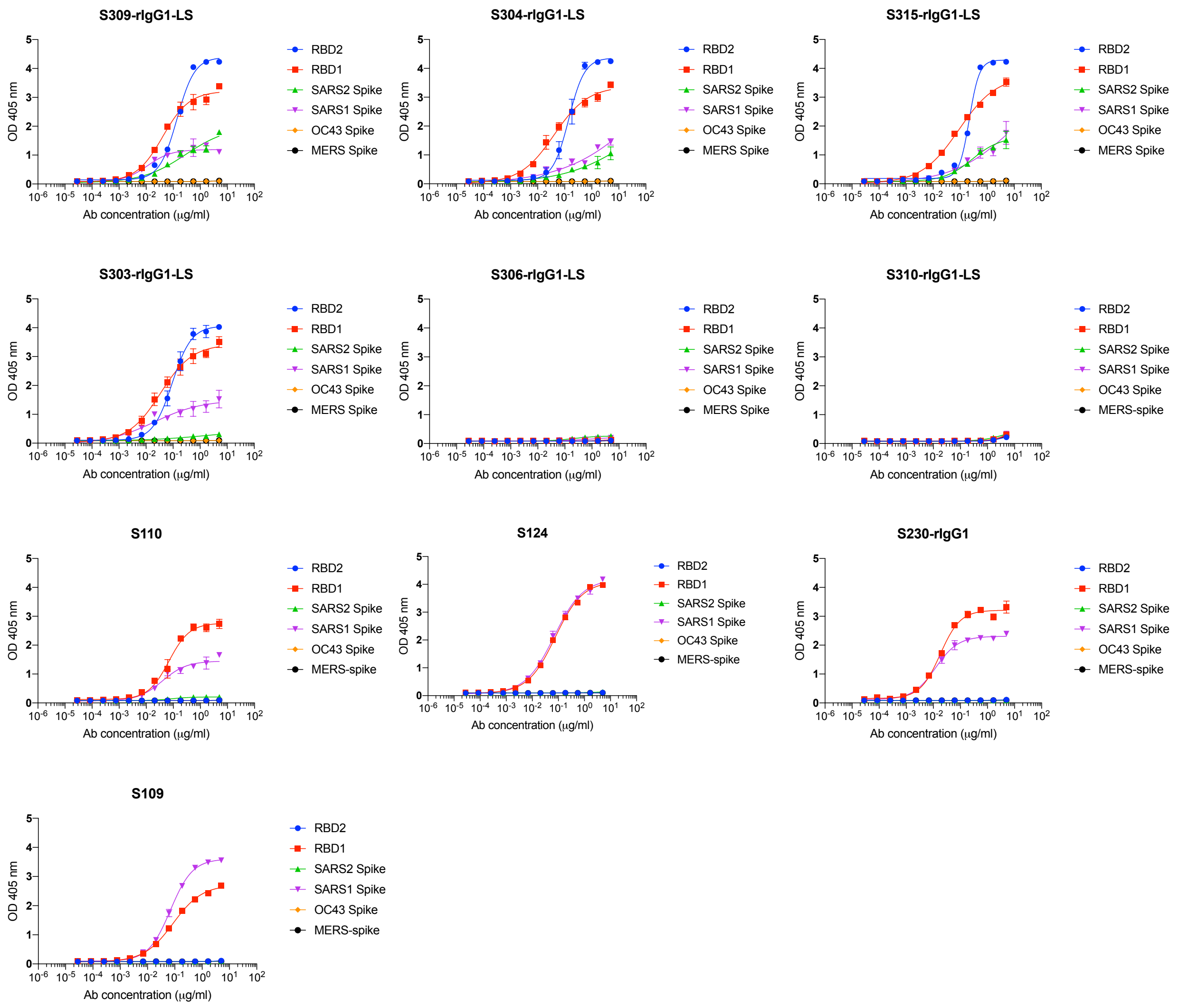


**Extended Data Fig. 1: Binding of cross-reactive antibodies to RBD of SARS-CoV (SARS1) and SARS-CoV-2 (SARS2) and to ectodomains of different coronavirus strains.** Recombinant mAbs were tested by ELISA at a concentration range of 5 to 0.00028μg/ml. RBD2: Receptor binding domain of SARS-CoV-2. RBD1: Receptor binding domain of SARS-CoV. Spike: stabilized prefusion trimer of the indicated coronavirus. Some antibodies were recombinantly expressed as IgG1 (rIgG1), some antibodies were recombinantly expressed as IgG1 with an LS mutation in the Fc part (rIgG1-LS).

**Extended Data Figure 1**

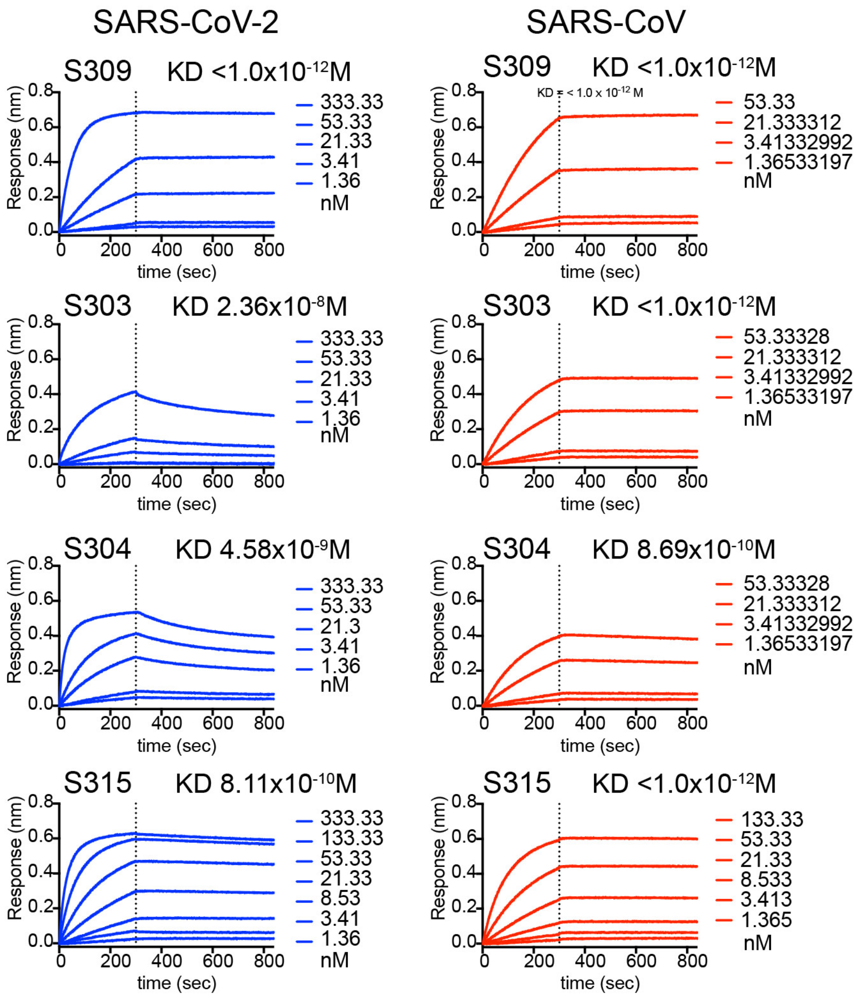


**Extended Data Fig. 2:** Antibody affinity/avidity of S309, S303, S304 and S315 to the RBD of SARS-CoV (red) and SARS-CoV-2 (blue). Antibodies were loaded to BLI pins via protein A for the measurement of association of different concentrations of RBD. Vertical dashed lines indicate the start of the dissociation phase when BLI pins were switched to buffer.

**Extended Data Figure 2**

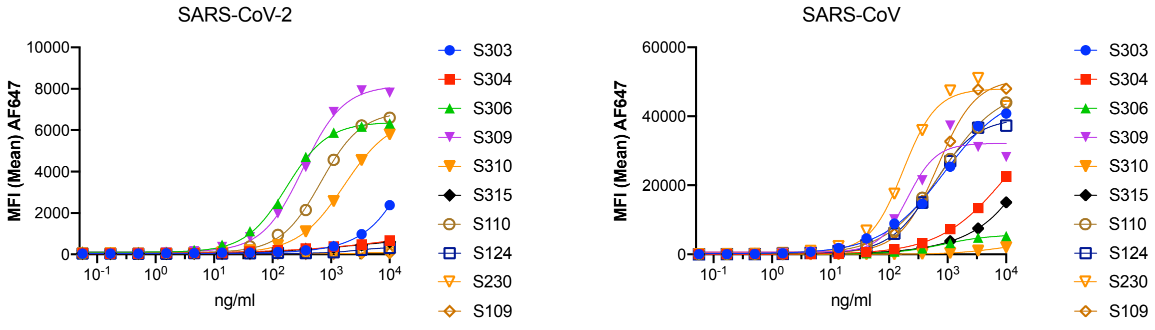


**Extended Data Fig. 3: Binding of crossreactive mAbs to ExpiCHO cells transfected with SARS-CoV S or SARS-CoV-2 S**. Mean fluorescence intensity as measured in flow cytometry for each antibody. Antibody concentrations tested are indicated in the x axis.

#### Extended Data Figure 3

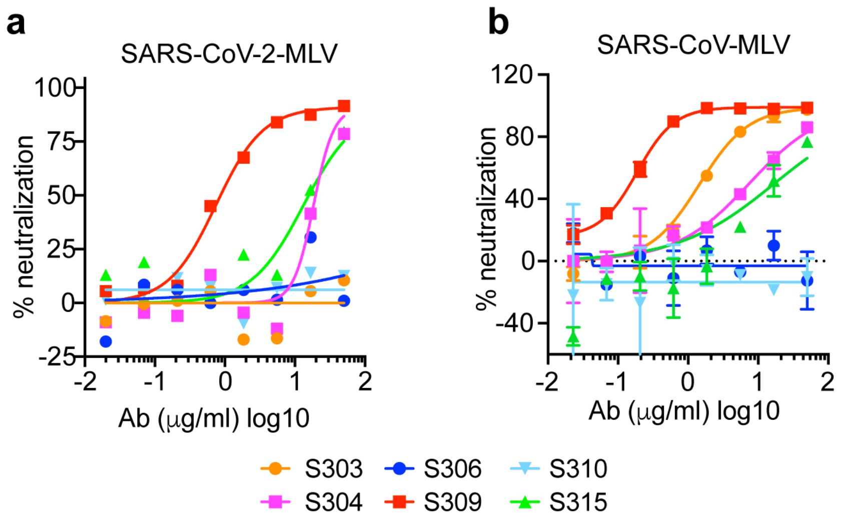


**Extended Data Fig. 4: Neutralization of cross-reactive antibodies. a-b,** the six cross-reactive mAbs indicated in the legend were tested for neutralization of SARS-CoV-2-MLV (a) and SARS-CoV-MLV (b).

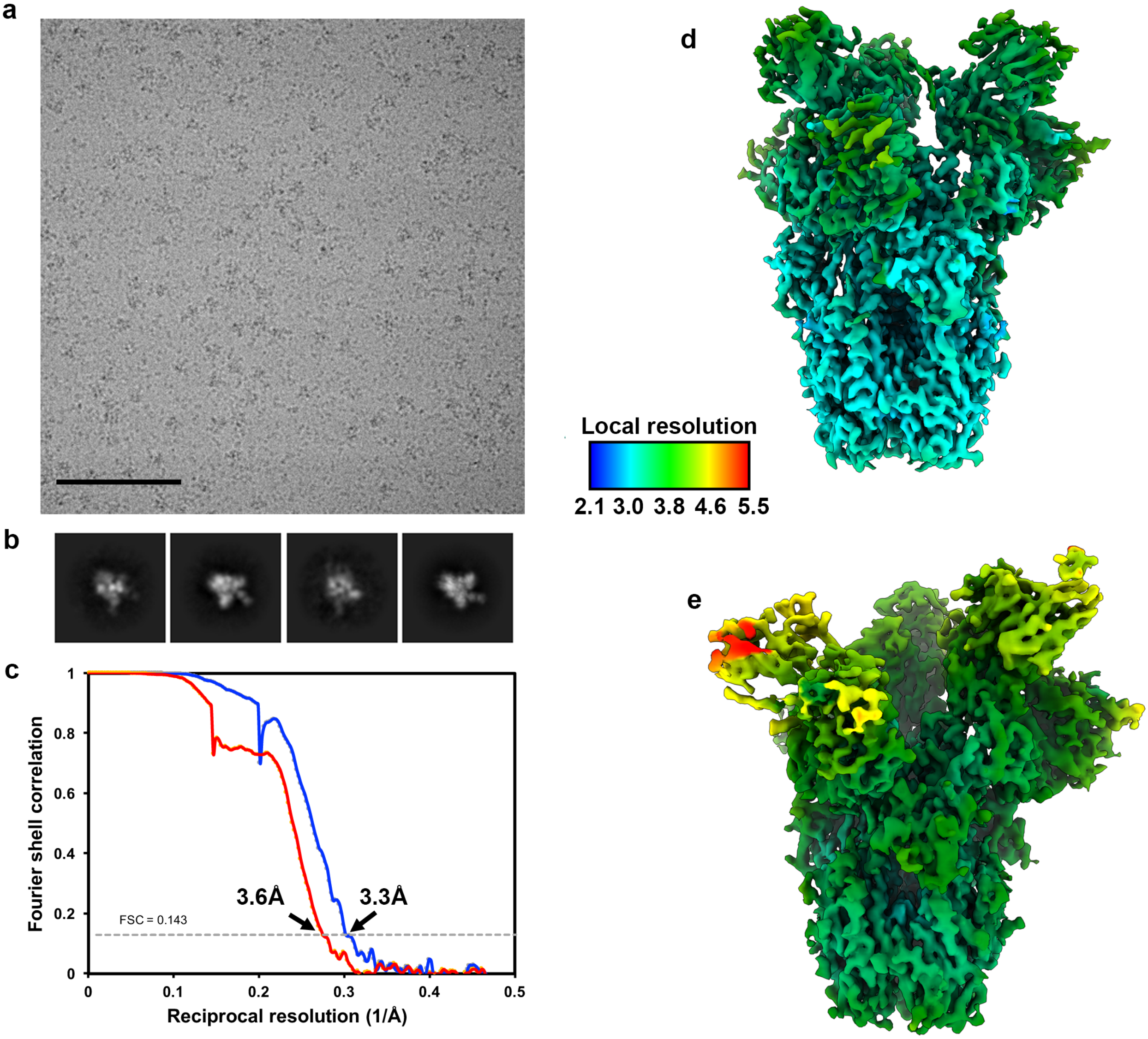


**Extended Data Fig. 5: CryoEM data processing and validation. a-b.** Representative electron micrograph (A) and class averages (B) of SARS-CoV-2 S embedded in vitreous ice. Scale bar: 100nm. **c.** Gold-standard Fourier shell correlation curves for the closed (blue) and partially open trimers (red). The 0.143 cutoff is indicated by horizontal dashed lines. **d-e.** Local resolution maps calculated using cryoSPARC for the closed (d) and open (e) reconstructions.

**Extended Data Figure 5**

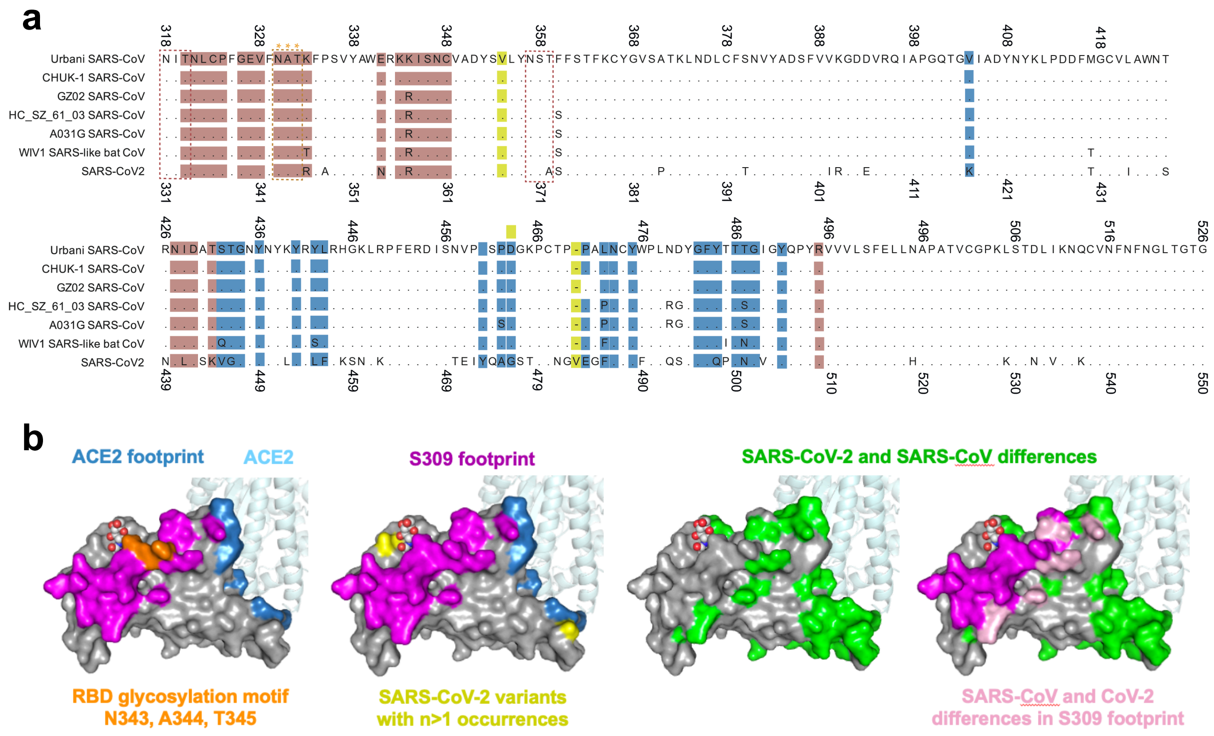


**Extended Data Fig. 6: Amino acid sequence conservation of the SARS-CoV-2 S$^{B}$ domain. a,** Alignment of the RBD of multiple sarbecoviruses. The ACE2 and S309 footprints are highlighted in blue and brown, respectively. The three high frequency variants are shown in yellow **b,** ACE2 and S309 footprints mapped onto the structure of the SARS-CoV-2 RBD domain (PDB ID: 6M0J). The ACE2 footprint was defined by residues being within 5Å of the receptor in 6M0J. The highly conserved NAT glycosylation motif is shown in orange (left), the SARS-CoV and SARS-CoV-2 differences within the S309 footprint is shown in pink (right), and the three high frequency RBD variants are shown in yellow (2$^{nd}$ from right).

**Extended Data Figure 6**

**a**

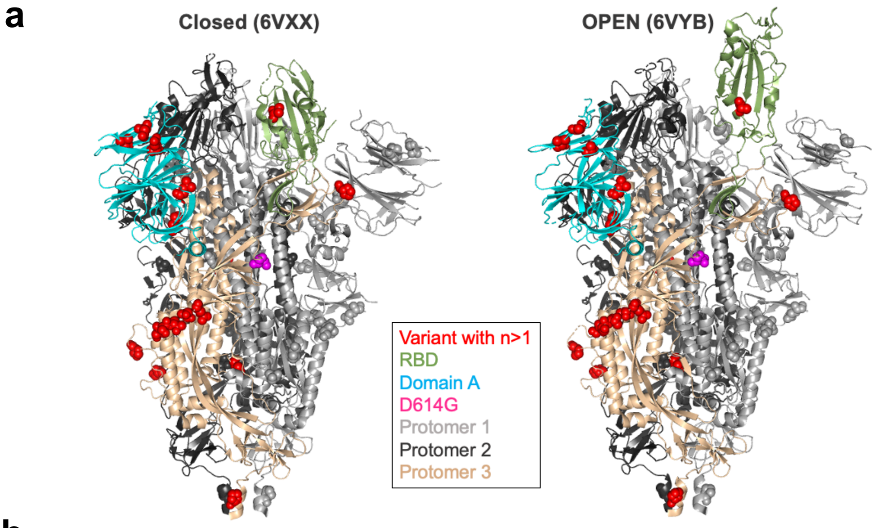


**b**

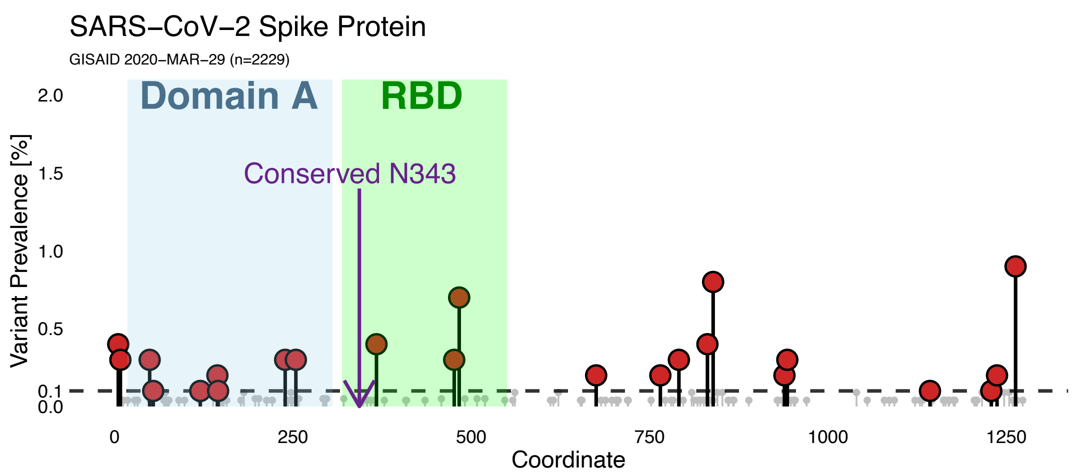


**c**

| | | SARS-CoV2 pos. | 333 | 334 | 335 | 336 | 337 | 339 | 340 | 341 | 343 | 344 | 345 | 346 | 354 | 356 | 357 | 358 | 359 | 360 | 361 | 440 | 441 | 442 | 444 | 509 | n | IC50 (µg/ml) |
|---|---|---|---|---|---|---|---|---|---|---|---|---|---|---|---|---|---|---|---|---|---|---|---|---|---|---|---|---|
| | | SARS-CoV pos. | 320 | 321 | 322 | 323 | 324 | 326 | 327 | 328 | 330 | 331 | 332 | 333 | 341 | 343 | 344 | 345 | 346 | 347 | 348 | 427 | 428 | 429 | 431 | 495 | | |
| Sarbecoviruses | Clade 2 | Human - SARS-CoV2 | T | N | L | C | P | G | E | V (I) | N | A (S) | T | R | N | K | R | I | S (N) | N | C | N | L | D | K | R | 2229 | 0.24 |
| | | GD Pangolin | - | - | - | - | - | - | - | - | - | - | - | T | - | - | - | - | - | - | - | - | - | - | - | - | 2 | |
| | | Bat CoV RaTG13 | - | - | - | - | - | - | - | - | - | - | - | T | - | - | - | - | - | - | - | H | I | - | - | - | | |
| | | GX Pangolin | - | - | - | - | - | - | - | - | - | - | S | K | - | - | - | - | - | - | - | K | Q | - | L | - | 6 | |
| | | Bat CoV ZC45, ZXC21 | - | - | V | - | - | H | K | - | - | - | - | - | E | T | K | - | - | D | - | K | Q | - | | - | 2 | |
| | Clade 3 | Human SARS-CoV | - | - | - | - | - | - | - | - | - | - | - | K | E | - | K / R | - | - | - | - | - | I | - | T | - | 53 | 0.12-0.15 |
| | | Civet SARS-CoV | - | - | - | - | - | - | - | - | - | - | - | K | E | - | - | - | - | - | - | - | I | - | T | - | | |
| | | Racoon dog SARS-CoV | - | - | - | - | - | - | - | - | - | - | - | K | E | - | - | - | - | - | - | - | I | - | T | - | | |
| | | Bat CoV WIV16 | - | - | - | - | - | - | - | - | - | - | - | - | E | - | - | - | - | - | - | - | I | - | T | - | | |
| | | Bat CoV WIV1 | - | - | - | - | - | - | - | - | - | - | - | T | E | - | - | - | - | - | - | - | I | - | T | - | | 0.18 |
| | | Other Bat CoVs | - | - | -/R | - | - | -/D | -/R/S (K/A) | -/I | - | - (V) | -/S | -/T | E | K/T (I/V) | -/K | - | - | -/D/E | - | K/Q (S) | H/Q/I (K) | - | G/T (S) | - | 21 | |
| | Clade 1 | Bat CoV BtKY72 | - | - | - | - | - | - | Q | - | - | - | S | N | E | L | - | - | - | D | - | S | V | - | - | - | | |
| | | Bat CoV BGR2008 | - | Q | - | - | - | N | - | - | - | I | - | S | E | M | - | - | T | - | - | S | - | - | | - | | |

() = variants found in only 1 sequence

**Extended Data Fig. 7: SARS-CoV-2 S residue conservation. a,** SARS-CoV-2 S protein variants occurring with a frequency of n>1 are shown as red spheres mapped onto the closed and open form of the full trimeric ectodomain. 40 mutations (out of 2229 total) are shown. Residues 476 (n=7) and 483 (n=17) are not shown since they were not resolved in the cryoEM structures. **b,** Prevalence of S variants throughout the amino acid sequence. Each dot is a distinct variant. The locations of $S^A$ and $S^B$ (RBD) are shown. Variants passing a frequency threshold of 0.1% (left) or 1% (right) are highlighted in red. Low-frequency variants (less than 2%, left) or all variants (right) are displayed. Position 614 is D in ≈60% and G in ≈40% of reported strains (not shown). **c,** Conservation analysis of S residues making contact with S309 across sarbecovirus clades, as defined in (Lu et al, Lancet 2020). Residue numbers for both SARS-CoV-2 and SARS-CoV are shown. The N343 NAT glycosylation motif is shown in orange. Sequence differences between SARS-CoV-2 and SARS-CoV are indicated in pink. n, number of sequences used for this analysis. IC50, mean half maximal neutralizing concentration of S309 against SARS-CoV or SARS-CoV-2-MLV.

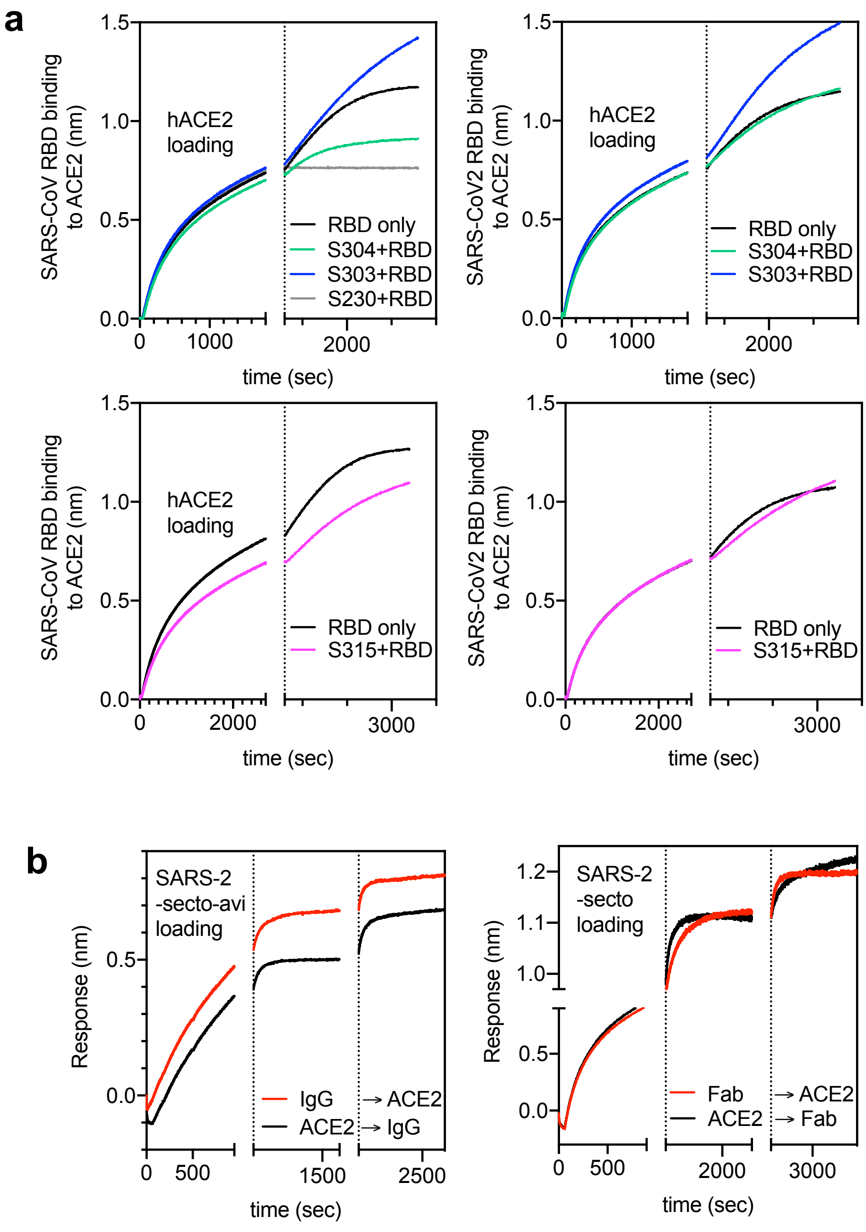


**Extended Data Fig. 8: Evaluation of competition between mAbs and ACE2 for binding to SARS-CoV-2 and SARS-CoV S glycoproteins. a,** Human ACE2 (hACE2) was loaded onto BLI sensors, followed by incubation of the sensors with RBD alone or RBD in combination with recombinant mAbs. The vertical dashed line indicates the start of the loading of RBD pre-complexed with mAbs or not. **b,** SARS-CoV-2 ectodomain was loaded onto BLI sensors, followed by incubation of the sensors with hACE2 or S309 IgG (left panel) or Fab (right panel). In a third step, sensors were incubated with hACE2 or S309 IgG/Fab as indicated.

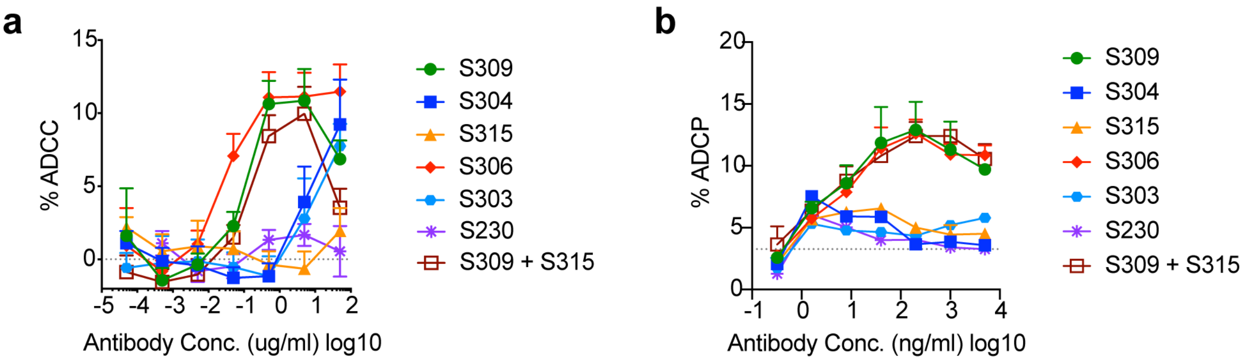


**Extended Data Fig. 9: ADCC and ADCP data for one representative donor. a,** ADCC for one donor homozygous for high affinity variant FcγRIIIa 158V (VV). Background signal of cells without mAbs was deducted from all values before plotting (dashed line indicates baseline). **b,** ADCP for one donor heterozygous for FcγRIIIa 158V (FV). The dashed line indicates the background signal for cells without mAbs.

**Extended Data Figure 9**

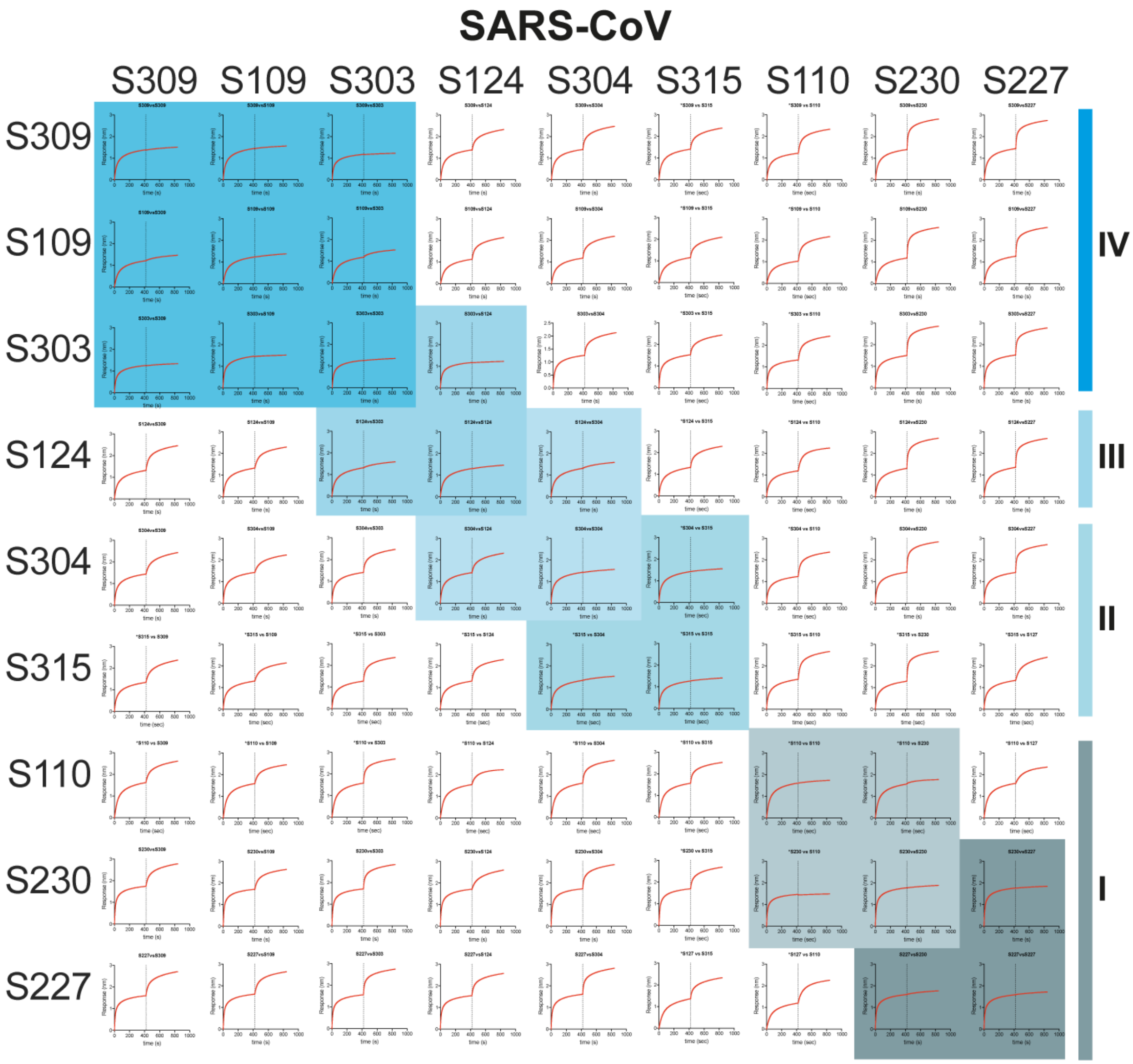


**Extended Data Fig. 10:** Competition of mAb pairs for binding to the RBD domain of SARS-CoV as determined by BLI (Octet). RBD was loaded on BLI pins. Association was measured first for antibodies indicated on the left of the matrix, followed by association of the antibodies indicated on top of the matrix.

### Extended Data Figure 10
